# Recovering genomic clusters of secondary metabolites from lakes: a Metagenomics 2.0 approach

**DOI:** 10.1101/183061

**Authors:** Rafael R. C. Cuadrat, Danny Ionescu, Alberto M. R. Davila, Hans-Peter Grossart

## Abstract

**Background:** Metagenomic approaches became increasingly popular in the past decades due to decreasing costs of DNA sequencing and bioinformatics development. So far, however, the recovery of long genes coding for secondary metabolism still represents a big challenge. Often, the quality of metagenome assemblies is poor, especially in environments with a high microbial diversity where sequence coverage is low and complexity of natural communities high. Recently, new and improved algorithms for binning environmental reads and contigs have been developed to overcome such limitations. Some of these algorithms use a similarity detection approach to classify the obtained reads into taxonomical units and to assemble draft genomes. This approach, however, is quite limited since it can classify exclusively sequences similar to those available (and well classified) in the databases.

In this work, we used draft genomes from Lake Stechlin, north-eastern Germany, recovered by MetaBat, an efficient binning tool that integrates empirical probabilistic distances of genome abundance, and tetranucleotide frequency for accurate metagenome binning. These genomes were screened for secondary metabolism genes, such as polyketide synthases (PKS) and non-ribosomal peptide synthases (NRPS), using the Anti-SMASH and NAPDOS workflows.

**Results:** With this approach we were able to identify 243 secondary metabolite clusters from 121 genomes recovered from the lake samples. A total of 18 NRPS, 19 PKS and 3 hybrid PKS/NRPS clusters were found. In addition, it was possible to predict the partial structure of several secondary metabolite clusters allowing for taxonomical classifications and phylogenetic inferences.

**Conclusions:** Our approach revealed a great potential to recover and study secondary metabolites genes from any aquatic ecosystem.

## Background

Metagenomics, also known as environmental genomics, describes the study of a microbial community without the need of *a priori* cultivation in the laboratory. It has the potential to explore uncultivable microorganisms by accessing and sequencing their nucleic acid [1]. In recent years, due to decreasing costs of DNA sequencing - metagenomic databases [2] (e.g., MG-RAST) have rapidly grown and archive billons of short read sequences [3]. Many metagenomic tools and pipelines were proposed to better analyse these enormous datasets [4]. Additionally, these tools allow to (i) infer ecological patterns, alfa- and beta-diversity and richness [5]; (ii) assemble environmental contigs from the reads [6] and more recently, (iii) recover draft genomes from metagenomic bins [7][8][9]. By recovering a high number of draft genomes from these so far uncultivable organisms, it is now possible to screen for new genes and clusters, unlocking a previously underestimated metabolic potential such as secondary metabolite gene clusters by using a metagenomic approach called Metagenomics 2.0 [10].

Polyketide synthases (PKS) and non-ribosomal peptide synthases (NRPS) are two families of modular mega-synthases, both are very important for the biotechnological and pharmaceutical industry due to their broad spectrum of products, spanning from antibiotics and antitumor drugs to food pigments. They act in an analogous way, producing polyketides (using acyil-coA monomers) and peptides (using aminoacyl monomers), respectively. Both families are broadly distributed in many taxonomical groups, ranging from bacteria (alphaproteobacteria, cyanobacteria, actinobacteria) to fungi [11][12].

PKS enzymes can be classified in types (I, II and III), where type I can be further classified into modular or iterative classes. The iterative PKS use the same domain many times, iteratively, to synthetize the polyketide. The modular PKS are large multi-domain enzymes in which each domain is used only once in the synthesis process [13][14]. The production of the polyketide follows the co-linearity rule, each module being responsible for the addition of one monomer to the growing chain [15].

Type I PKS are characterized by multiple domains in the same open reading frame (ORF) while in type II each domain is encoded in a separate ORF, acting interactively [16]. Type III is also known as Chalcone synthase and has different evolutionary origin from type I and II [17]. Type III PKSs are self-contained enzymes that form homodimers. Their single active site in each monomer catalyzes the priming, extension, and cyclization reactions iteratively to form polyketide products [17]. Hybrid PKS/NRPS and NRPS/PKS are also modular enzymes, encoding lipopeptides (hybrid between polyketides and peptides) and occur in bacterial as well as fungal genomes [18][19][20].

PKS and NRPS are very well explored in genomes from cultivable organisms, mainly *Actinomycetes* [21] and *Cyanobacteria* [22]. Recently, by using a metagenomic approach, studies have demonstrated the presence of these metabolite-genes in aquatic environments, as for example, Brazilian coast waters (in free living and particle-associated bacteria) and from the microbiomes of Australian marine sponges [23]. However, there are few metagenomic studies whose scope is to find these gene families in freshwater environments where most studies are based on isolation approaches [24][25].

In addition, due to the rather large size of genes involved in these pathways, yet, it is not possible to recover the full genes by using traditional read-based metagenomics or the single sample assembly approach. Most of the studies aim to solely find specific domains, like Keto-synthase (KS) in PKS and Condensation domain (C) in NRPS, due to the high conservation of these domains [26].

We used a metagenomics 2.0 approach to overcome these limitations and improve the screening for secondary metabolism genes and clusters while evaluating the potential of microbial communities for future research on potential drugs. This study aims to (i) generate draft genomes from Lake Stechlin; (ii) to screen these genomes for new complete multi-modular enzymes from PKS and NRPS families, exploring their diversity and phylogeny.

## Methods

### Sampling and sequencing

A total of 26 metagenomic samples from Lake Stechilin, north-eastern Germany were used. Water was collected as metagenomic samples on several occasions (April, June 2013, July 2014, Aug 2015) in sterile 2 L Schott bottles from Lake Stechlin (53°9’5.59N, 13°1’34.22E). All samples, except those from Aug 2015, were filtered through 5 μm and subsequently 0.2 μm pore-size filters. The samples collected in Aug 2015 were not size-fractionated and directly filtered on a 0.2 μm pore size filter due to specific research demands. Genomic DNA was extracted using a phenol/chloroform protocol as described in [27] and was sent for sequencing.

Sequencing was conducted at MrDNA (Shallowater, Texas) on an Illumina Hiseq 2500, using the V3 chemistry, following, fragmentation, adaptor ligation and amplification of 50 ng genomic DNA from each sample, using the Nextera DNA Sample Preparation Kit.

Table S1 shows the general information about the 26 samples used in this study.

### Environmental draft genomes

Briefly, all samples were pre-processed by Nesoni (https://github.com/Victorian-Bioinformatics-Consortium/nesoni) to remove low quality sequences and to trim adaptors, and afterwards assembled together using MegaHIT (default parameters) [6]. The reads from each sample were mapped back to these assembled contigs using BBMAP (https://sourceforge.net/projects/bbmap/) and then all data was binned using MetaBAT [7] to generate the draft genomes. The completeness and taxonomical classification were checked using CheckM [28].

### Screening secondary metabolism genes and phylogenetic analysis of NRPS and PKS domains

DNA fasta files of the generated bins (288) were submitted to a locally installed version of Anti-SMASH (–clusterblast –smcogs –limit 1500) [29]. Using in-house ruby scripts, the domains from PKS and NRPS were parsed. The PKS KS domains and NRPS C domains were submitted to NAPDOS for classification [30]. In addition, all the KS and C domains (trimmed by NAPDOS) were submitted to BLASTP against RefSeq database [31], using the default parameters. The 3 best hits of each domain were extracted and added to the original multi-fasta file with the environmental domains. The full set of KS and C domains (from bins and references obtained by the blast on RefSeq database) was submitted for NAPDOS for the phylogenetic analysis. The resulting alignment and tree were exported and the trees were manually checked and annotated.

### Relative abundance of bins in each sample

The reads from each sample were mapped (using BBMAP) against each bin fasta file and an in-house ruby parser script was used to calculate the relative abundance of each bin in each sample, normalizing the read counts by the number of reads of each sample. The table with the results was loaded into STAMP [32] in order to analyse the significant differences of bin abundance over the samples.

## Results

### Environmental draft genomes obtained (bins)

Metagenomic binning resulted in 288 draft environmental genomes (called bins in this study). Of these, 45 had a predicted completion level of more than 75% according to CheckM.

Table S2 shows the general information about each bin, including completeness, genome size, number of open reading frames (ORFs) and taxonomical classifications (from CheckM).

### Screening secondary metabolism genes and phylogenetic analysis

By using Anti-SMASH, at least one secondary metabolite gene cluster was found in 121 of the bins, totalling 243 clusters and 2200 ORFs. From these 243 clusters, 125 (51.4%) were classified in the Terpene and 35 (14.40%) in the Bacteriocin pathway. In addition, a total of 18 NRPS, 6 type I PKS and 3 hybrid PKS/NRPS clusters were found in 15 different bins (Figure 1a). The latest 3 obtained pathway clusters are the main focus of our study.

Figure 1b shows the taxonomical classification at phylum level for the bins showing NRPS, type I PKS and hybrid clusters. Supplementary table S3 shows the distribution of all clusters in all bins.

**Figure 1A:**
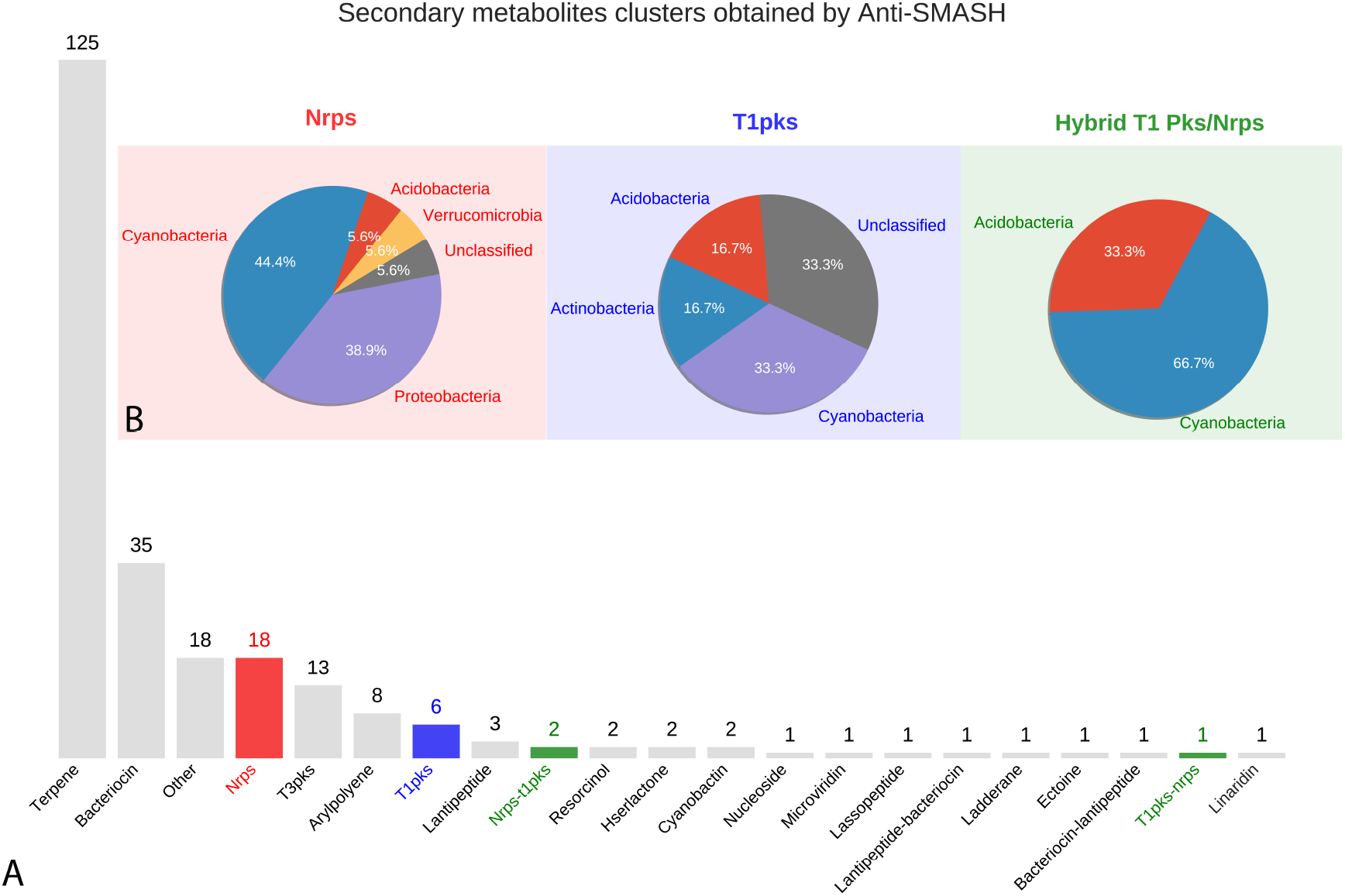
Abundance of secondary metabolite cluster types obtained with Anti-SMASH in the recovered 288 bins (environmental genomes). B: Taxonomical classification of bins (Phyla) in which NRPS, PKS and Hybrid PKS/NRPS clusters were found. Red bar and pie: NRPS; blue bar and pie: Type I PKS; green bars and pie: Hybrid clusters (NRPS-PKS and PKS-NRPS).

A total of 43 condensation (C) domains were obtained from NRPS clusters. All these sequences were submitted to NAPDOS analysis. Figure 2a shows the classification of C domains into classes. Most of the sequences were classified as LCL domains (58%). This kind of domain catalyzes the formation of a peptide bond between two L-amino acids.

**Figure 2A:**
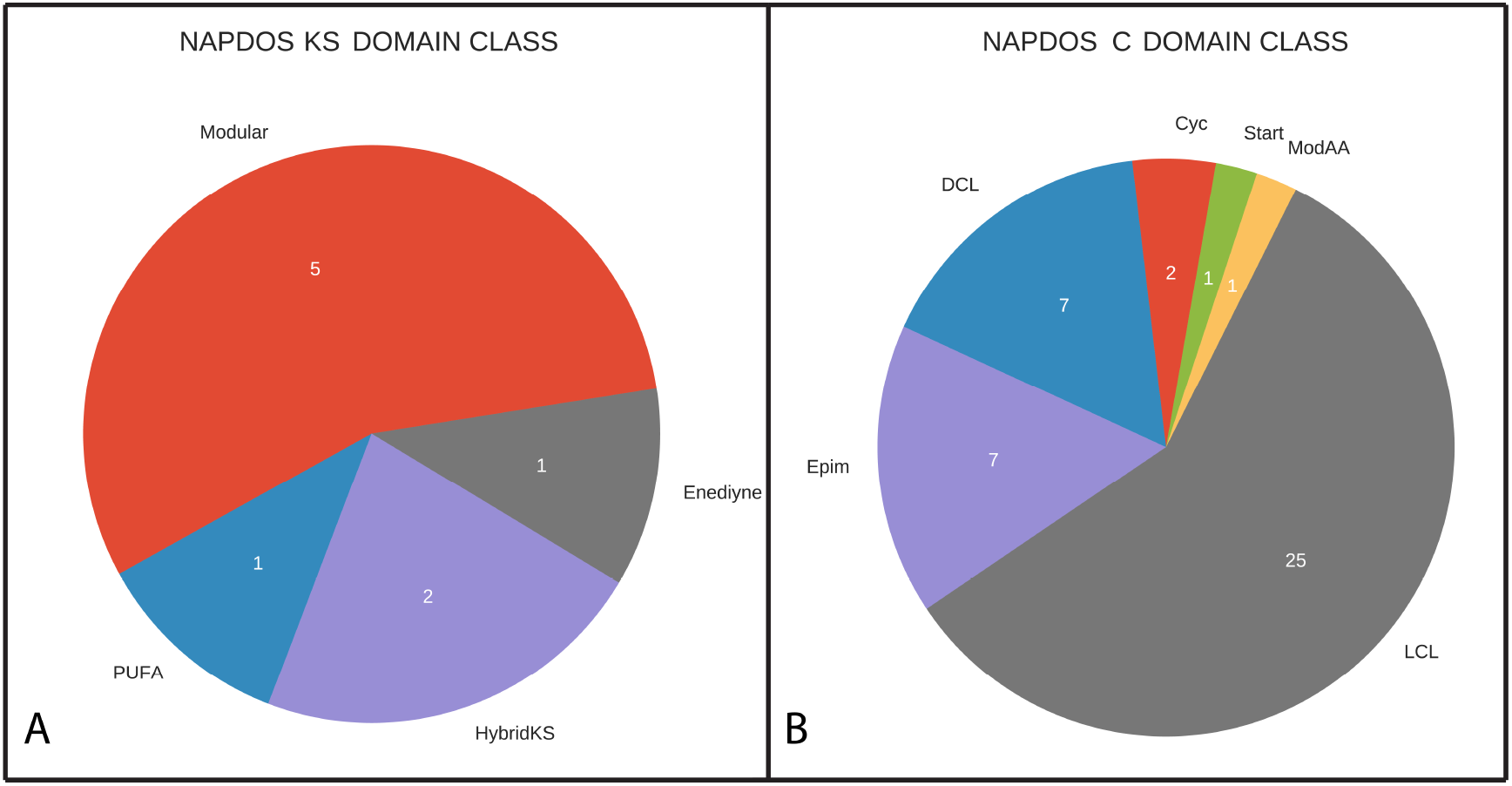
NAPDOS classification of the NRPS KS domain. Modular: possess a multidomain architecture consisting of multiple sets of modules; hybridKS: are biosynthetic assembly lines that include both PKS and NRPS components; PUFA: Polyunsaturated fatty acids (PUFAs) are long chain fatty acids containing more than one double bond, including omega-3-and omega-6-fatty acids; Enediyne: a family of biologically active natural products. The Enediyne core consists of two acetylenic groups conjugated to a double bond or an incipient double bond within a nine-or ten-membered ring. **2B: NAPDOS classification of NRPS C domain.** Cyc: cyclization domains catalyze both peptide bond formation and subsequent cyclization of cysteine, serine or threonine residues; DCL: link an L-amino acid to a growing peptide ending with a D-amino acid; Epim: epimerization domains change the chirality of the last amino acid in the chain from L-to D-amino acid; LCL: catalyze formation of a peptide bond between two L-amino acids; modAA: appear to be involved in the modification of the incorporated amino acid; Start: first module of a Non-ribosomal peptide synthase (NRPS).

The screening for type I PKS resulted in 9 KS domain sequences. Most of them are classified as modular type I PKS (56%). All of them were submitted to NAPDOS and classified into 4 different classes (Figure 2b).

All the KS and C domains were also submitted to similarity analysis by using BLASTP against RefSeq database (table S4 and S5), and the best 3 hits of each sequence were extracted and used for phylogenetic analyses with NAPDOS. The trees for C and KS domains are shown in Figures 3 and 4, respectively.

**Figure 3:**
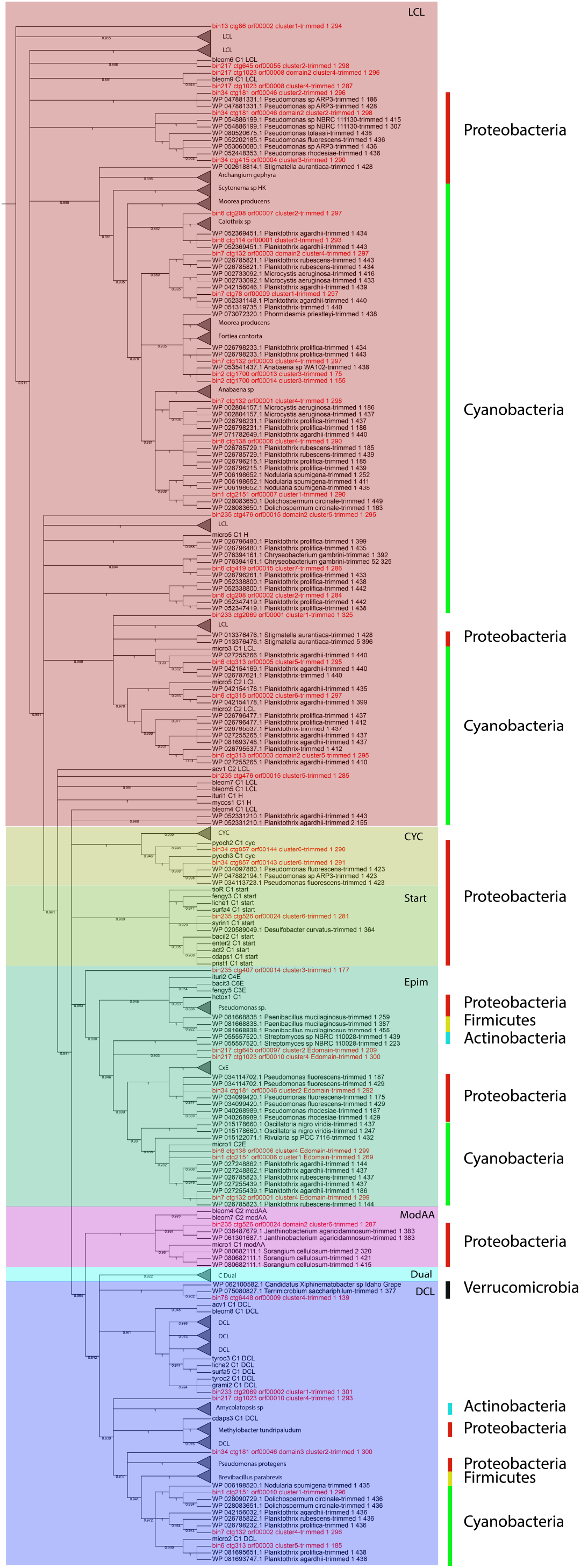
NAPDOS phylogenetic tree of C domains (environmental domains, the top 3 blast results on RefSeq and the NAPDOS reference sequences). The shadow colours represent the domain classifications (LCL, CYC, Start domains, EPIM, ModAA, Dual, and DCL). The sidebars represent phyla (Proteobacteria, Cyanobacteria, Firmicutes, Actinobacteria, Verrucomicrobia). All sequences from environmental bins are in red.

**Figure 4:**
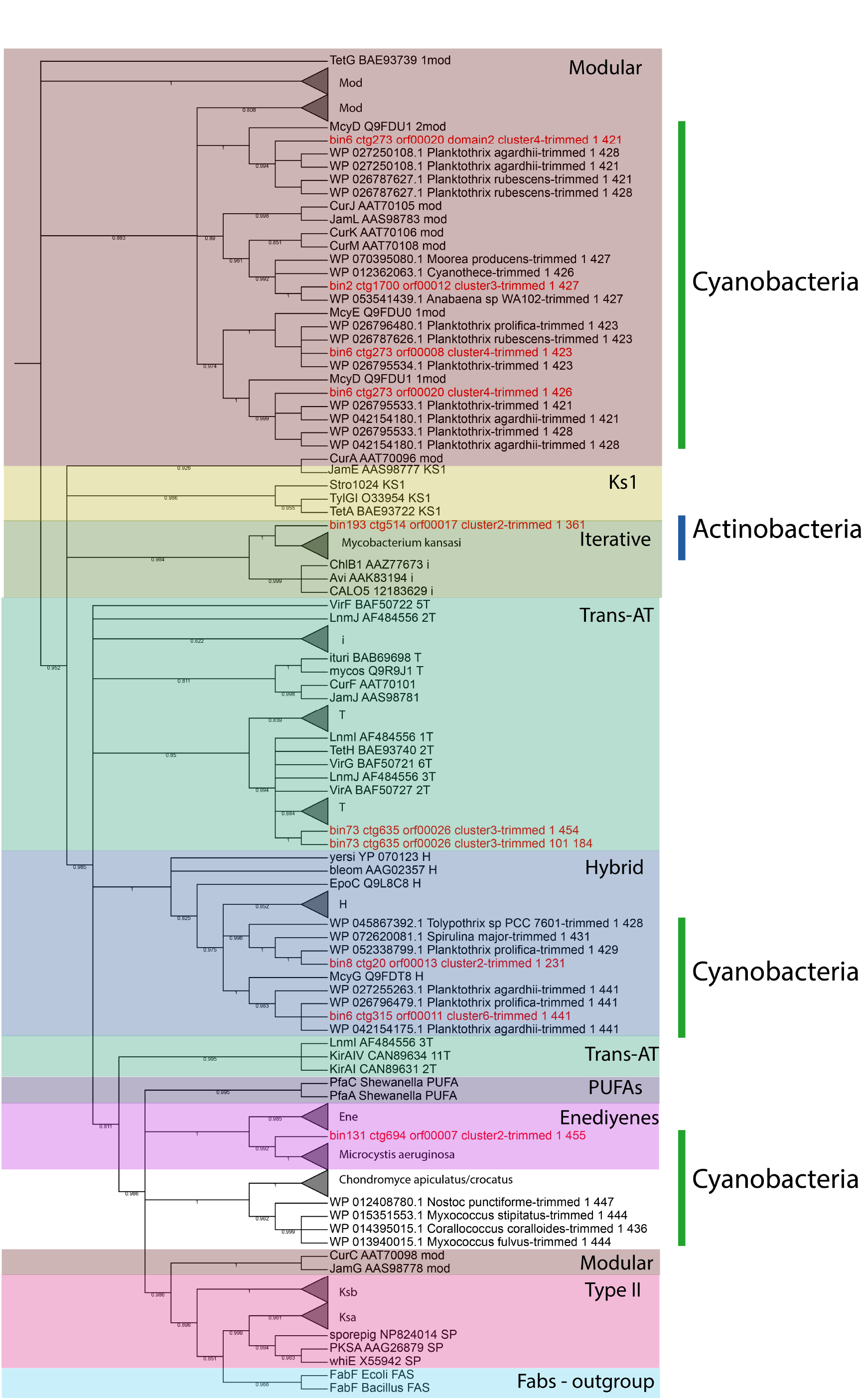
NAPDOS tree of KS domains (environmental domains, the top 3 blast results on RefSeq and the NAPDOS reference sequences). The shadow colours represent the domain classifications (Modular, KS1, Iterative, Trans-AT, Hybrid, PUFA, Enediyenes, Type II and Fabs - Fatty acid synthase). The sidebars represent phyla (Cyanobacteria and Actinobacteria). All the sequences from environmental bins are in red.

### Relative abundance of bins on each sample

The relative abundance of all bins in each sample was estimated by mapping the reads from each sample against the assembled bins. Table S6 shows the normalized bin abundance for every sample.

Due to differences between the filtration methods, we decided to classify the samples in 3 groups: particle associated samples (PA) – filtered on 5 μm membranes (and also samples from aggregates), free-living samples (FL) – pre-filtered through 5.0 μm membranes and subsequently filtered on 0.22 μm membranes, and non-size fractionated samples (NSF) - filtered direct on 0.22 μm membranes (without previous filtering) retaining the whole bacterial community. In total, we obtained 7 samples in the PA group, 5 in the FL group, and 14 in the NSF group.

The table with the relative abundance of the bins in all samples was loaded on STAMP and an ANOVA test was conducted followed Games-Howell POST-HOC test and Benjamini-Hochberg FDR correction. Table S7 shows 158 bins for which the difference in relative abundance was statistically significant (p < 0.05) between the 3 groups (FL, PA and NSF).

From the 15 bins containing NRPS and/or type I PKS clusters, only 4 showed significant difference between the 3 groups. Bins 1 and 2 are more abundant in PA samples and bins 193 and 235 are more abundant in the FL samples.

### Exploring NRPS, type I PKS and hybrid clusters from draft genome bins

We highlight 3 bins (with less than 35% contamination and more than 70% completeness) out of the 15 obtained type I PKS and/or NRPS and explore their clusters.

In bin 34 (*Pseudomonas*, 98.28% completeness) it was possible to retrieve 7 clusters, including 3 NRPS clusters (Figure 7) and 2 Bacteriocin clusters.

In cluster 2 (ctg181), multiple domains of NRPS (with the 3 minimal modules) and regulatory genes were identified, e.g., smCOG: SMCOG1057 (TetR family transcriptional regulator) (Figure 5, in green arrows).

**Figure 5:**
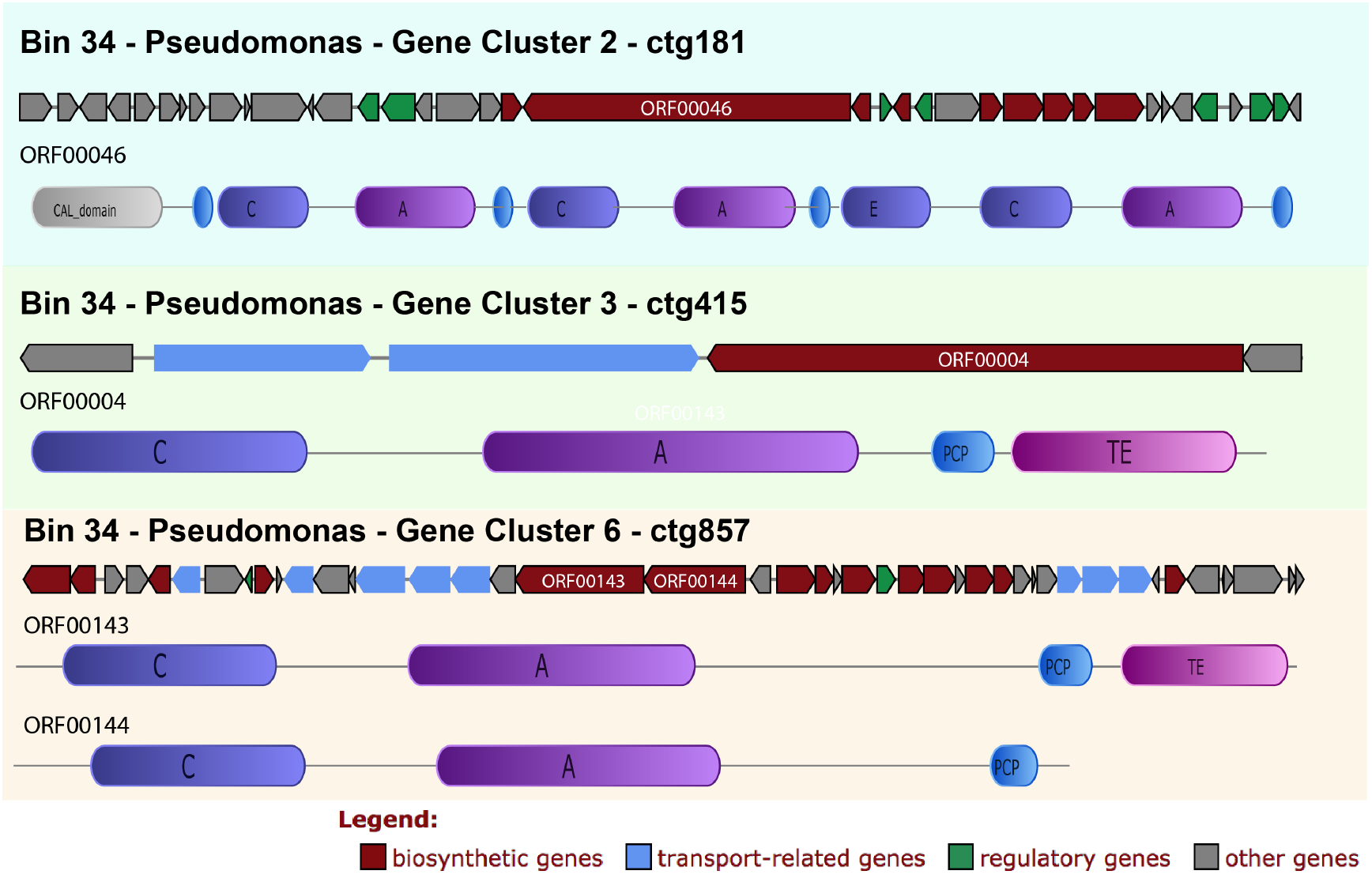
Bin 34, NRPS clusters detailed annotation and synteny. The synteny of the clusters with a functional classification for each ORF is given. In addition, for the NPRS biosynthetic ORFS the domain annotations are given. CAL: Co-enzyme A ligase domain, C: condensation, A: adenylation, E: epimerization, TE: Termination, KR: Ketoreductase domain, and ECH: Enoyl-CoA hydratase

All clusters show a high similarity with *Pseudomonas* proteins. Cluster 2 has a similarity of 92% with *Pseudomonas synxantha* bg33r, conserving also the gene synteny.

The C domain sequences were submitted to NAPDOS analysis and 2 were classified as belonging to the Syringomycin pathway and the LCL class, and one was classified as belonging to the Microcystin pathway or/and the DCL class (links an L-amino acid to a growing peptide ending with a D-amino acid).

In cluster 3 (ctg415) (Figure 5), in addition to the NRPS domains, the following transporter related genes are found: smCOG: SMCOG1288 (ABC transporter related protein) and SMC0G1051 (TonB-dependent siderophore receptor) (blue narrows). Nevertheless, this cluster is not complete and just one C domain was found (LCL class), which was also classified to the Syringomycin pathway.

Cluster 6 (ctg857 – Figure 5) shows many NRPS domains, regulatory factors and transporters genes, including drug resistance genes, e.g. SMC0G1005 (drug resistance transporter, EmrB/QacA), SMC0G1044 (ABC transporter, permease protein) and SMC0G1051 (TonB-dependent siderophore receptor) (blue arrows). Two C domains from this cluster were classified as belonging to the heterocyclization class. This class catalyzes both peptide bond formation and subsequent cyclization of cysteine, serine or threonine residues [33]. Both domains were classified in the Pyochelin pathway by NAPDOS. The phylogenetic tree (Figure 3) confirms both the functional and taxonomical classification (confidence value 100).

**Figure 6:**
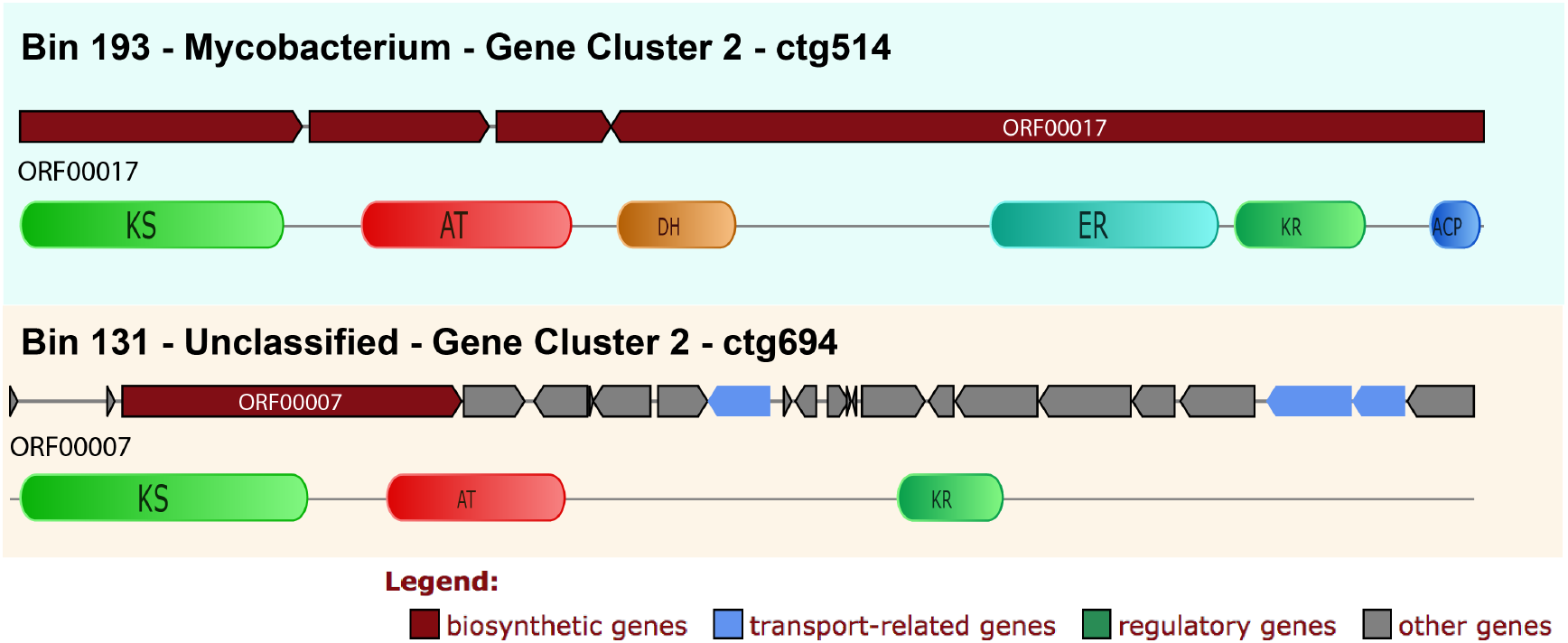
Bin 193 and 131 type I PKS clusters detailed annotation and synteny. It is possible to see the synteny of the cluster with the functional classification for each ORF. In addition, for the PKS biosynthetic ORF the domain-specific annotations can be seen. KS: keto-synthase, AT: acyltransferase, KR: ketoreductase, E: epimerization, DH: eehydratase, and ER: enoylreductase.

In **bin 193** (Mycobacterium, 73.37 % completeness) a type I PKS cluster was identified (ctg514) (Figure 6). Five PKS domains were retrieved, including the minimal core from one of the ORFs on this contig. The KS domain BLASTP result shows 82% (and 99% coverage) similarity with *Mycobacterium kansasii*. The NAPDOS analysis from the KS domain suggests that it could be a modular (Epothilone pathway) or iterative type I PKS similar to the Calicheamicin pathway. However, by using the phylogenetic analysis it was clustered as the iterative clade (confidence value 98.4) together with the *Mycobacterium kansasii* sequence (confidence value 100) (Figure 4).

Two further clusters were recovered: one type III PKS and one unclassified one. All clusters show similarity with the *Mycobacterium* clusters.

Bin 131 (unclassified bacteria by CheckM) has 84.09% of completeness reported by CheckM. In this bin it was possible to find one cluster and 3 domains of type I PKS (KS, AT and KR) (Figure 6). The KS domain was classified by NAPDOS as belonging to the Maduropeptin and Neocarzinostatin pathways. Using clusterblast inside Anti-Smash it was not possible to find any similar cluster, but using BLASTP it was possible to find similarity with the cyanobacteria *Microcystis aeruginosa* (64% identity and 99% coverage on BLASTP search). In the phylogenetic tree, it was clustered within the Enediynes clade (Figure 4) and also with *Microcystis aeruginosa* (confidence value 99).

In addition, there are 12 more bins with NRPS or type I PKS clusters, but with less than 70% of completeness or more than 35% contamination. The bins 1 (69.54% completeness) and 2 (16.52%) were classified as the genus *Anabaena*, showing NRPS and hybrid NRPS-type 1 PKS, respectively. Bins 6 (39.66% completeness), 7 and 8 were classified as the genus *Planktothrix* and show a high diversity of secondary metabolites: 3 NRPS clusters, one type I PKS and 2 NRPS-PKS hybrid (bin 6), 2 NRPS (bin 7) and 2 NRPS and one type I PKS (bin 8). The bin 8 also shows a Microviridin cluster.

Bins 73 and 217 are classified as *Acidobacteria* showing PKS and NRPS, respectively. Bin 235 (*Burkholderiaceae* family) shows 3 NRPS clusters and bin 13 (*Comamonadaceae* family, also from *Burkholderiales* order) shows one NRPS cluster.

Bin 78 is classified as *Verrucomicrobiaceae* shows 1 NRPS cluster. Additionally, there is an unclassified *Archaea* (NRPS cluster) and one bin without any classification (bin 136, type I PKS).

The Anti-Smash results for all bins are available in the Supplemental Information (SI 1).

## Discussion

The field of metagenomics has generated a vast amount of data in the last decades [34]. Most of the data is poorly annotated and little quality controlled when loaded into the public databases, hence awaiting a more in-depth analysis [35]. There are many open challenges in this field, e.g., (i) the lack of representative genomic databases from uncultivable organisms to be used in a similarity-based annotation procedure; (ii) high confidence assembly of short reads from species-rich samples; (iii) obtaining high enough coverage for every organism in the sample, including those with a low abundance, etc. [36]. Recently, some new algorithms have been proposed to overcome these limitations and to obtain partial or near complete genomes from environmental samples, e.g., MetaBat [7] and MetaWatt [8]. Most of them require many samples and high coverage sequencing per sample as an input. Recently, studies have been done to recover genomes even from rare bacteria [37]. The term Metagenomics 2.0 was introduced to describe this new generation of metagenomic analysis by Katherine McMahon [10] and most of the studies using this approach have been conducted to reveal ecological interactions and networks [38][39].

In this study, we recovered 288 environmental draft genomes using 26 samples from Lake Stechlin, a temperate oligo-mesotrophic lake. One of the advantages of this approach is to enable the recovery of large genomic clusters, especially the Megasynthases clusters of the secondary metabolism, e.g., involved in biosynthesis of antibiotics, including its regulatory and transporter genes. Here, we have used the Anti-SMASH and NAPDOS pipelines to identify, annotate, classify and to carry out the phylogenetic analysis of a total of 243 clusters of known secondary metabolites. To our knowledge, this is the first study using the metagenomics 2.0 approach to recover Megasynthases clusters. A number of previous studies had been conducted using a traditional PCR based screening [40] and shotgun metagenomics approach [41][42] exploring the abundance and diversity of individual genes and domains, but these studies are missing the genomic context. By obtaining the entire genomic context it is possible, in future studies, to clone and to do heterologous expression for all the genes, including promoters and transporters.

Screening the bins for secondary metabolite clusters, we can see that the most abundant cluster belongs to the Terpene pathway (125 clusters) (Figure 1). This biosynthesis pathway is well known to be present in many plant and fungi genomes, but recently it was proposed to be also widely distributed in bacterial genomes. One study revealed 262 distinct terpene synthases in the bacterial domain of life [43]. Consequently, it can represent a fertile source of new natural products – yet, greatly underestimated. The second most abundant class of clusters belongs to the Bacteriocin pathway (35 clusters) (Figure 1). Bacteriocin is a group of ribosomal synthesized antimicrobial peptides which can kill or inhibit bacterial strains closely related or non-related to the Bacteriocin producing bacteria [44]. It has been suggested as a viable alternative to traditional antibiotics and can be used as narrow-spectrum antibiotics [45]. Only a few studies have been conducted to screen for Bacteriocin genes by using a metatenomic approach, and solely for the host-associated microbiome [46] or fermented food microbiome [47][48][49]. None of these studies was conducted for natural environments and none have used the metagenomics 2.0 approach.

In this study, we focused on 2 families of large modular secondary metabolite genes, type I PKS and NRPS. With our approach, it was possible to find a total of 18 NRPS, 6 type I PKS and 3 hybrid PKS/NRPS clusters. For NRPS clusters, it was possible to recover 43 C domains, most of them (58%) from the LCL class. An LCL domain catalyzes a peptide bond between two L-amino acids [50]. A previous study also found that the LCL class was the most abundant in another aquatic environment, dominated by gram-negative bacteria [42]. Many studies have shown that the LCL class in aquatic environments is limited to gram-negative bacteria [51]. Our results further support this as we also found the LCL class only in bins of gram-negative bacteria (Figure 3) with the only exception of the unclassified Archaea (bin 233), which should be further investigated in order to confirm the phylogenetic classification. It was also possible to recover 9 KS domains from the type I PKS clusters, 56% from the modular class and 22% from the hybrid PKS/NRPS class. Those classes are larger (with many copies of each domain) than the iterative ones, increasing the chances to be recovered by metagenomic approaches. Accordingly, the NRPS and PKS clusters were more in depth analyzed, including syntheny, domain phylogeny, and partial metabolite protein structure predictions.

From the bins showing secondary metabolite genes, the most complete was from bin 34 (*Pseudomonas*). The 7 clusters on this genome vary from 8,675 base pairs (bp) to 52,516 bp in size, been only possible to be recovered due to the high completeness of the assembled genome (98.28%). The presence of a great diversity of clusters in Pseudomonas is expected, as many active secondary metabolites (encoded by NRPS) have been previously described in the Pseudomonas genus, ranging from antibiotics and antifungal to siderophores [52][53][54].

From those 7 clusters in bin 34, the NRPS clusters 2 and 3 showed a high similarity with Syringomycin (three domains) and Microcystin (one domain) pathways. The first one is found, for example, in the *Pseudomonas syringae* (a plant pathogen) genome, as a virulence factor (Syringomycin E) [55] which also has antifungal activity against *Saccharomyces cerevisiae* [56]. On the other hand, Microcystin is a class of toxins produced by freshwater Cyanobacteria species [57] and it can be produced in large quantities during massive bloom events [58]. Due to the taxonomical classification of the bin and the higher number of domains similar to Syringomycin, however, it is more likely that the product encoded by this cluster is functionally close to the latter pathway.

In bin 34 - cluster 6, both C domains were classified as belonging to the Pyochelin pathway. This peptide is a siderophore of *Pseudomonas aeruginosa* [59]. The presence of a TonB-dependent siderophore receptor in cluster 6 provides additional evidence about its functional classification. Additionally, two Bacteriocin clusters and one Aryl polyene cluster were found in bin 34. Aryl polyenes are structurally similar to the well-known carotenoids with respect to their polyene systems and it was recently demonstrated that it can protect bacteria from reactive oxygen species, similarly to what is known for carotenoids [60]. These results suggest that a wide range of metabolites is encoded in this *Pseudomonas* genome, providing it an “arsenal” of secondary products, increasing the likelihood of the *Pseudomonas* species to succeed in aquatic systems.

Bin 131 (unclassified bacteria) shows a PKS cluster and 3 domains. It was classified as belonging to the Enediynes pathway. These compounds are toxic to DNA and are under investigation as anti-tumor agents, with several compounds under clinical trials [61]. All are encoded by type I iterative PKS [62] and it was possible to recover the minimal core (KS, AT and KR) as well as transporter genes from the environmental genome. The most similar PKS I present in the public databases stems from *Microcystis aeruginosa*, but only with an identity of 64%, suggesting that it is encoding for a new compound, which has not previously been described.

In bin 193 (*Mycobacterium* - sister linage *M. rhodesiae*), one of the 3 recovered clusters is a Type I PKS, similar to iterative PKS in the NAPDOS analysis (confirmed by the phylogenetic analysis). The most similar KS sequence belongs to *Mycobacterium kansasii*, with 82% similarity. *M. rhodesiae* and *M. kansasii* are both non-tuberculous mycobacteria (NTM) that can be found in different environments, but both can also be opportunistic pathogens and cause a chronic pulmonary infection in immunosuppressed patients [63]. The species *M. kansasii* comprises various subtypes and some are often recovered from tap water and occasionally from river or lake water [64][65]. There is still controversy about how the transmission from environment to human host occurs and also about the implications on public health. The presence of PKS in Mycobacterium genus was discovered more than a decade ago and most of the polyketides encoded by different species of this genus play role in virulence and/or components of the extraordinarily complex mycobacterial cell envelope [66]. Further studies must be done in order to investigate better the potential of this bin to cause infections on humans, i.e. by screening virulence factors on the full genome.

Five bins of the phyla Cyanobacteria contained PKS and NRPS clusters. In bins 1 and 2 (Anabaena, now called “Dolichospermum”), it was possible to recover 2 NRPS and 1 hybrid NRPS-PKS, respectively. The genus *Anabaena* is known to encode several toxins, including the dangerous Anatoxin-a, and to produce toxic blooms in lakes and reservoirs [67][68][69][70]. However, the Anatoxin-a is encoded by a type I PKS cluster [71], unlike the NRPS and Hybrid clusters found in the *Anabaena* bins from this study.

On the other hand, the hepatotoxic heptapeptide of the class Mycrocystin is present in many genera of Cyanobacteria, including *Anabaena*, and they are encoded by NRPS and also Hybrid NRPS-PKS clusters [72][73][74][75]. The results of NAPDOS reveal one C domain from bin 1 classified in the pathway of Mycrocystin with e-value 6e-83.

In bins 6, 7 and 8 (*Planktothrix*), it was possible to find several type I PKS, NRPS and hybrid clusters. In bin 6 there are 3 NRPS, one type 1 PKS and one hybrid cluster. The bin 7 shows 2 NRPS and bin 8 shows one type I PKS and 2 NRPS clusters. The genus *Planktothrix* also can be producer of Anatoxin-a [75] and the presence of type I PKS cluster on these bins can be alarming. However, the 3 KS domains from type I PKS from bin 6 reveal great similarity with the Epothilone pathway. The NapDOS analysis and the KS domain from bin 8 suggest a high similarity to the neurotoxin Jamaicamides pathway. A previous study showed in 2010 [76] the presence of an Anatoxin-a-producing cyanobacterium in northeastern Germany Lake Stolpsee, rising concerns about the presence of these toxins in the waters of these lakes.

The absence of Anatoxin-a genes in the studied lake is in agreement with previous screening for a toxin screening in the lake [77].

In the bin 73 (*Acidobacteriales*) 1 PKS sequence was found. By the phylogenetic classification, its KS domain is clustered with *trans*-AT KS domains. The AT domains of *trans*-AT PKSs are not integrated into the assembly lines but expressed as free-standing polypeptides, unlike the more familiar *cis*-AT PKSs [78]. However, the NAPDOS result shows the AT domain of this bin in the same ORF with KS and KR domains, showing a syntheny that suggests a *cis*-AT PKS. In addition, the classification by similarity from NAPDOS suggests a polyunsaturated fatty acid (PUFA) but only with 31% of identity

To assess the life style of the bins (free-living or particle-associated), we calculated the relative abundance of the bins in every sample. A total of 158 bins with significant difference between the 3 groups were found (Table S7), however from the 15 bins on which this study focused (showing NRPS and/or type I PKS clusters), only 4 bins (26.6%) were significantly differently present in the life-styles. Bins 1 and 2 (*Anabaena* genus) are more abundant in the PA group, especially on samples B7 and B9 (and also the replicates Old_b7 and Old_b9), accounting for 20-25% (bin 1) and 10-15% (bin 2) on these samples. The very high abundance of these bins on the samples can be explained, based on the long term monitoring program of IGB on Lake Stechlin, by the fact that these samples were collected during the occurrence of a massive cyanobacterial bloom.

From the other bins containing PKS/NRPS clusters, we can see that Bins 6, 7, and 8 (*Planktothrix*), beside the lack of significant difference between FL and PA groups (p-value > p 0.05), they are clearly more abundant in NSF. The possible explanation for this notion is that the NSF samples were collected during a mesocosm experiment, whereas the other samples were directly derived from the lake.

## Conclusions

Using the Metagenomics 2.0 approach, we were able to recover full megasynthases sequences and its genomic context from environmental draft genomes. However, there are limitations, e.g., the genomic coverage of less abundant organisms and the possibility of chimeras. Recently, it has been demonstrated that with an increasing number of samples, it will be possible to recover individual species genomes with a high confidence [7]. In the near future, with the advent of the 3^rd^ generation sequencing, with longer reads, up to 100 kilobases, it will be possible to further improve the quality of the assemblies [79]. These new approaches unlock the possibility of studying these newly recovered environmental pathways and their evolution in detail. Thus, allowing cloning and expressing these clusters will provide new insights on natural products of great interest for biotechnological and pharmaceutical industry. Moreover, studies have demonstrated the possibility to synthesise large functional DNA [80], and together with additional screening techniques, it will be possible to obtain such sequences and to synthesise the full cluster for heterologous expression, skipping the cloning and functional screening process, saving considerable time and money. In addition, the current work highlights the great potential for the discovery of new metabolically active compounds in freshwaters such as oligo-mesotrophic Lake Stechlin. Further, the study of complete or near complete genomes from uncultivated bacteria in the natural environment will enable us to better understand the multiple forms of interactions between species and how they compete for the limiting natural resources.

## Declarations

### Available of data and materials

The sequences generated for this study (metagenomic reads) were deposited in ENA (PRJEB22274 and PRJEB7963).

### Authors’ contributions

Conceived and designed the experiments: RRCC, DI, AMRD, and HPG. Performed the experiments: RRCC, DI. Analyzed the data: RRCC, DI, and HPG. Contributed reagents/materials/analysis tools: DI and HPG. All authors wrote the manuscript and revised it for significant intellectual content.

## Acknowledgements

We thank Dr. Camila Mazzoni and all the team of Berlin Center for Genomics in Biodiversity Research (BeGenDiv) for allowing us to use the facilities and computational resources for the bioinformatics analyses. Elke Mach and the MIBI group are thanked for their technical support and fruitful discussions.

